# An NMDA-R mediated short-term memory resistant to anesthesia in adult *Danio rerio*

**DOI:** 10.1101/795351

**Authors:** Gokul Rajan, Joby Joseph

## Abstract

Memory in animals is labile in the early phase post-training. Memory in the early phase has been shown to be disrupted by treatments such as electroconvulsive shock or cold-shock (Quinn and Dudai, 1976). Using hypothermic shock and other pharmacological interventions, the various underlying memory pathways in *Drosophila* can be identified as an immediate short-lasting anesthesia sensitive memory and a delayed anesthesia resistant memory which is followed by a more stable protein synthesis dependent long-term memory (Margulies *et al*., 2005). In another ectothermic animal, *Danio rerio*, a popular vertebrate model, we ask if such a memory component exists which is sensitive to hypothermic disruption. To test this, we developed a fear conditioning assay with a green light at the bottom of the tank as the conditioned stimulus (CS) and electric shock as the unconditioned stimulus (US). We also standardized a cold anesthesia protocol in adult zebrafish to induce stage V anesthesia. The learning/memory was found to be NMDA-R mediated. Cold anesthesia as well as tricaine mediated anesthesia did not significantly affect the early-acting memory trace induced by a fear-conditioning protocol in adult zebrafish. We suggest future directions to tease out the underlying memory components in the early phase of memory in zebrafish.

## INTRODUCTION

The development of our understanding of memory formation and consolidation was significantly influenced by the early works in invertebrate organisms like *Drosophila, Aplysia* and others. Memory retention in animals is known to be disrupted if electroconvulsive shock or anesthesia is applied shortly after training (Quinn and Dudai, 1976).

Cold-shock anesthesia administered by subjecting a fly in a test tube to 0° C ice water reduced the 3 hour memory retention in the flies. This effect became progressively less when the time delay between training and cold-shock anesthesia was increased (Tully *et al*., 1994). This was called the Anesthesia Sensitive Memory (ASM) in *Drosophila*. Although the loss in 3 hour memory retention was the highest when the delay between training and cold-anesthesia treatment was the least, the effect vanished beyond a delay of approximately half an hour (likely following an anesthesia resistant pathway). There was no effect of cold-shock aesthesia when it was given before the training.

Tully *et al*. (1994) showed that the memory that persisted in *Drosophila* 30 minutes post-training in the absence of any anesthesia consisted of two genetically and functionally independent components: anesthesia-resistant memory (ARM) and long-term memory (LTM). However, LTM was disrupted by cycloheximide, a protein synthesis inhibitor whereas ARM was resistant to cycloheximide treatment. A Drosophila mutant *radish* was also identified which was deficit in ARM; the LTM component remained unaffected in these mutants. (Folkers *et al*., 1993 and Tully *et al*., 1994).

Besides *radish* mutant, using an odor-avoidance response in *Drosophila*, Tully and Quinn (1985) identified *dunce* (PDE analog) and *rutabaga* (Ca-Calmodulin dependent Adenyly Cyclase) mutants deficient in early component of the memory (showing rapid decay in the initial 30 minutes); the later memory components remained intact. Protein synthesis dependent LTM component of the memory showed defects in many mutants like dBREB2, Adf1, Notch, crammer and nebula (Margulies *et al*., 2005).

Our understanding of the ASM and ARM pathways at a molecular and circuit level has increased in the recent times. At the molecular level, we know that *rutabaga* encodes calmodulin-dependent adenylyl cyclase (Levin *et al*., 1992). On the other hand, *radish* encodes a protein product which has phosphorylation sequences for PKA and can interact with a GTPase that regulates cytoskeleton rearrangement and hence, it may play a role in changing synaptic morphology (Folkers, 2006). This may also suggest that both ASM and ARM converge at cAMP/PKA signaling pathway. More recent works have highlighted the role of Synapsin and Bruchpilot in ASM and ARM respectively (Knapek *et al*., 2010 and Knapek *et al*., 2011). At the neuronal circuit level, it is suggested that ASM and ARM pathways are represented simultaneously in different subsets of Kenyon cells (Knapek *et al*., 2011).

Despite this understanding, it is far from clear why a cold-shock anesthesia would lead to a *rutabaga*-like deficit in short-term memory (STM). We hypothesize that in an ectothermic animal like *Drosophila*, a cold-shock anesthesia may temporarily arrest metabolic activities in a non-specific manner. This can show more pronounced effects on the temporally fast acting ASM pathway contributing to STM (than the slow-acting ARM pathway contributing to LTM). Hence, the observed deficit in STM caused by cold shock. Here we ask whether such a phenomenon would be reproducible in another ectothermic but vertebrate animal, *Danio rerio*.

*Danio rerio* is increasingly becoming a model of choice to study stress, anxiety, social behavior, learning and memory, in addition to its classical use in developmental biology and biomedicine (Bonan and Norton, 2015, Gerlach and Lawrence, 2008). There are many learning paradigms established in larval and adult zebrafish. These have shown the importance of NMDA-R and protein synthesis in associative memory (Blank *et al*., 2009 and Hinz *et al*., 2013). In the current work, we first developed a fear conditioning assay and a cold-shock anesthesia protocol for adult zebrafish. Later, we tested the effect of cold-shock anesthesia post-training on the early phase of memory retention to comment on the ASM pathway in adult Zebrafish.

## METHODS

### Animal procurement and housing

Adult zebrafish (*Danio rerio*) were used in the experiments. The adult fish (>5 weeks old) were bought from Sunny Aquariums, Hafeezpet, Hyderabad (and occasionally from other neighboring aquariums). Once procured, they were maintained in home tanks at ~28°C with 12 hour light and 12 hour dark cycle, for at least a week before performing any behavioral experiment. Fish were fed dried bloodworms. All procedures were approved by the institutional animal ethics committee.

### Training set-up

The fear conditioning assay was carried out in a custom built glass tank (18cm × 6cm × 12cm) with 200 ml water. The training set-up was also custom built. It comprised of two light lamps placed beneath two halves of the tank and a 9V battery connected to parallel steel mesh along the inside walls of the tank. The two lamps and the 9V battery were connected to three separate relays controlled by a microcontroller (ATtiny85) which was loaded with the training protocol (written on Arduino).

### Fear conditioning assay

A massed training protocol was used for classical conditioning assay. The assay was adapted and modified from Huang *et al* (2012). A green light at the bottom of the tank was used as the only conditioned stimulus (CS), while the unconditioned stimulus (US) was a 9V shock. The lit half of the tank was maintained at a light intensity of 152.33 +/− 3.38 lux and the unlit half was maintained at 9.33 +/− 1.20 lux. The home tanks are maintained at 48 +/− 1.53 lux. In the training mode, there is a 10 minutes baseline period and a 4 minutes test period, before and after the training period respectively. During the first half of the baseline and test phase, the CS randomly turns on either halves of the tank and then switches the position in the next half of the phase. A simultaneous-conditioning paradigm was used in the training phase where the CS and US turn on simultaneously. The massed training protocol lasts for 1.5 minutes with 9 training sessions. Each training session consists of 1.5 seconds of CS delivered with shock. The inter-session gap is 9.5 seconds. In the test mode, the total protocol runs for 10 minutes. The CS randomly turns on in one half of the tank and later flips over after 5 minutes. See supplementary video (S1.avi) for an example.

Video segments of 1 or 2 mins from each phase of the protocol were selected and split into 1fps images for manual analysis. The performance index (PI) of the fish was calculated after manually scoring the fish based on its position in the tank. The animal receives a score of +1 or −1 based on whether it is present in the non-CS area or CS area respectively in a given image. The PI is the average of the scores in a given time window selected for analysis. A random mix of male and female fish were used in the experiments to avoid any sexual bias affecting the results.

### Pharmacological assay

To block NMDA receptors a sub-anesthetic concentration (2 mg/L) of ketamine which is not known to cause any severe behavioral changes in zebrafish was used as a non-competitive glutamatergic antagonist in the study (Riehl *et al*., 2011). The fish was treated with 2mg/L ketamine for 20 minutes (10 minutes prior to beginning the training protocol and for another 10 minutes during the baseline period). The assay tank contained 2 mg/L ketamine. The training and scoring was done as described before. The PI of untreated control group and the ketamine treated test group was compared using unpaired t-test.

### Cold anesthesia protocol

The protocol involved transferring the fish into a Petriplate. The temperature of the Petriplate water was then lowered by keeping it at 0° C for approximately 8 minutes until a stage V anesthesia is obtained. The fish was then transferred to home tank maintained at room temperature (28 deg Celcius). The delay in transferring the fish from the training set-up to cold treatment ranged from 2 to 4 minutes.

Swimming behavior of fish treated with cold anesthesia were recorded at the following time intervals: baseline (before cold treatment), 0 minute (immediately after cold treatment), 30 minutes, and 60 minutes. The animal’s swimming movements in the recordings were tracked using NSYSLAB Tracker (animal tracking tool developed in our lab). From the positional coordinates, the following parameters were quantified using custom written scripts in MATLAB: mean velocity, initial latency to leave the lower one third of the tank, total time spent in lower third of the tank, cumulative duration spent in the central part of the tank and frequency of zone transition between the lower third and the rest of the tank. Control group consisted of fish which were similarly processed in a Petridish at room temperature. Comparisons between test group and control group were carried out for each parameter using unpaired t-test.

### Effect of cold anesthesia on memory

Fish in the control group and test group were maintained in equivalent conditions and processed alternately during the training and testing phase. After the training session, the test group animals would immediately undergo the aforementioned cold anesthesia treatment at 0° C whereas the control group animals would be sham treated at room temperature. The animals were then maintained in their home tanks for 2 hours. Now, the animals were tested in the same shuttle box task and scored for their performance as described before. The performance in this task is a measure of their memory retention. Comparisons between the two groups were carried out using unpaired t-test.

### Chemical anesthesia protocol

Tricaine solution (100mg/L) was used for chemical anesthesia. One fish at a time was transferred to the solution for 15 minutes. It took approximately 6.3 (+/− 1) mins for the fish to loose equilibrium and then the fish was maintained in this solution until the end of the 15 minutes. Once removed from the solution, it took 35.57 (+/− 4.10) secs for the fish to recover and start swimming again. The delay in transferring the fish from the training set-up to anesthesia treatment ranged from 2 to 4 minutes.

## RESULTS

### Massed training induced memory in *Danio rerio* is robust

**Figure 1:**
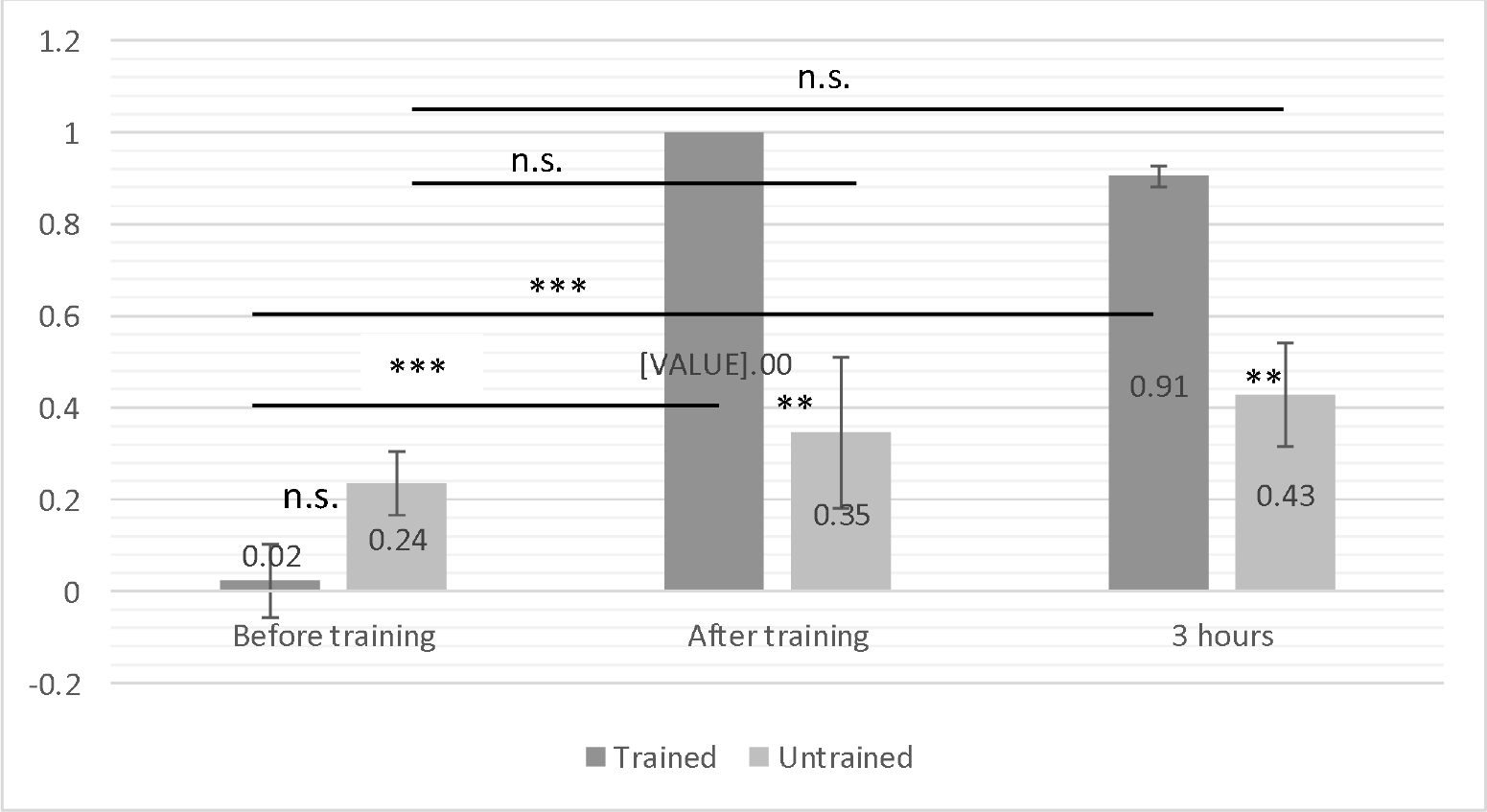
Performance index (PI) (mean+/−SE) in trained and untrained groups. The trained group (n=6) was trained on the custom built training protocol. An untrained control group (n=5) also went through the same protocol but without the US (9V shock) component. The performance of the two groups in the memory task was evaluated immediately after training and 3 hours post training. The test group shows a significant improvement in PI immediately after training (p<0.001) and 3 hours after training (p<0.001). In the untrained control group no significant increase in performance was noted. To determine the degree of significance, paired t-tests were performed for comparisons within the trained and untrained groups whereas unpaired t-tests were performed for comparisons between the trained and untrained group. ***p<0.001, **p<0.01

### Massed training induces a short-lived memory trace

**Figure 2:**
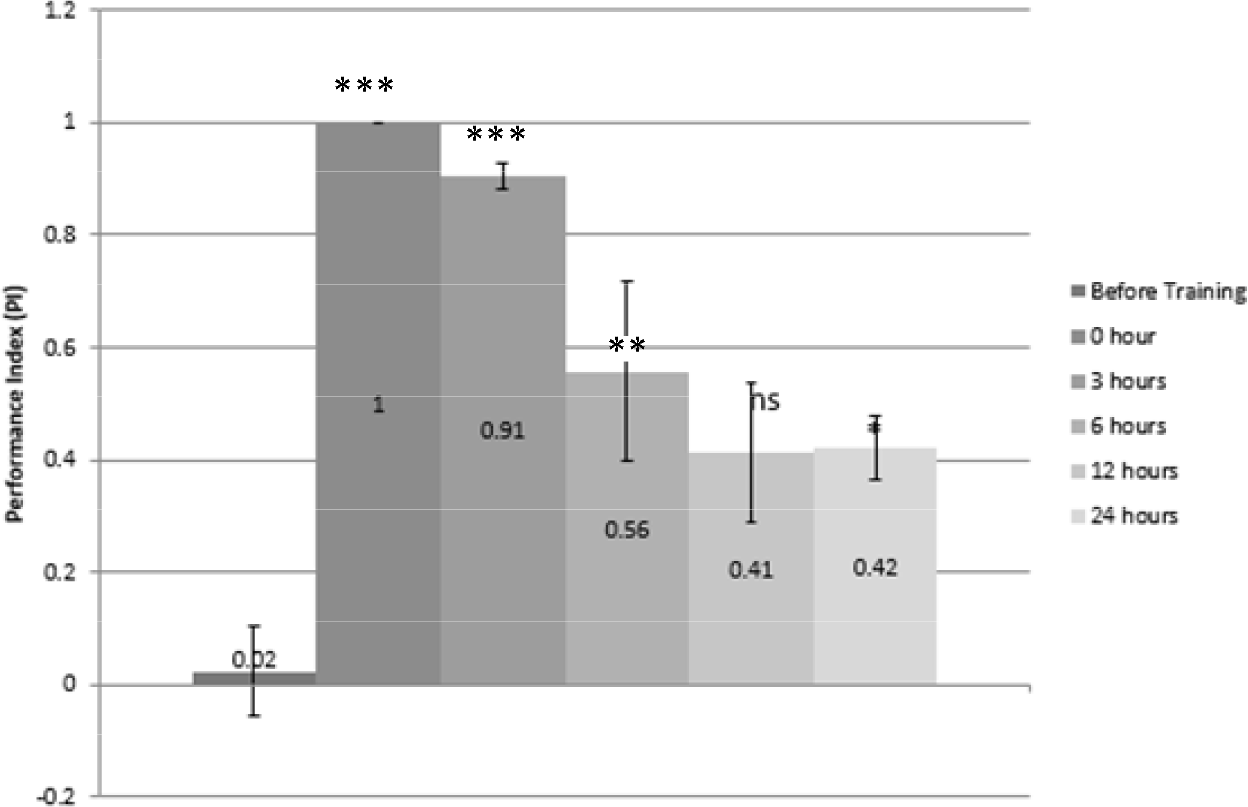
Decline in PI (mean+/−SE) of trained fish group over time. The performance of the trained group (n=5) remained high with high significance (p<0.001) for upto 3 hours post-training. At the 6^th^ hour, the PI was still significantly higher when compared to the pre-training behavior (p<0.01). At 12 hours post-training, the increase in PI was non-significant and at 24 hours post-training, the increase in PI was only weakly significant (p=0.037). This shows how the massed training protocol induces a short-lasting memory for few hours in contrast to a long lasting memory which stays robust for days. Paired t-tests were performed to determine the degree of significance. ***p<0.001, **p<0.01, *p<0.05

### Learning/recall is NMDA-R mediated

**Figure 3:**
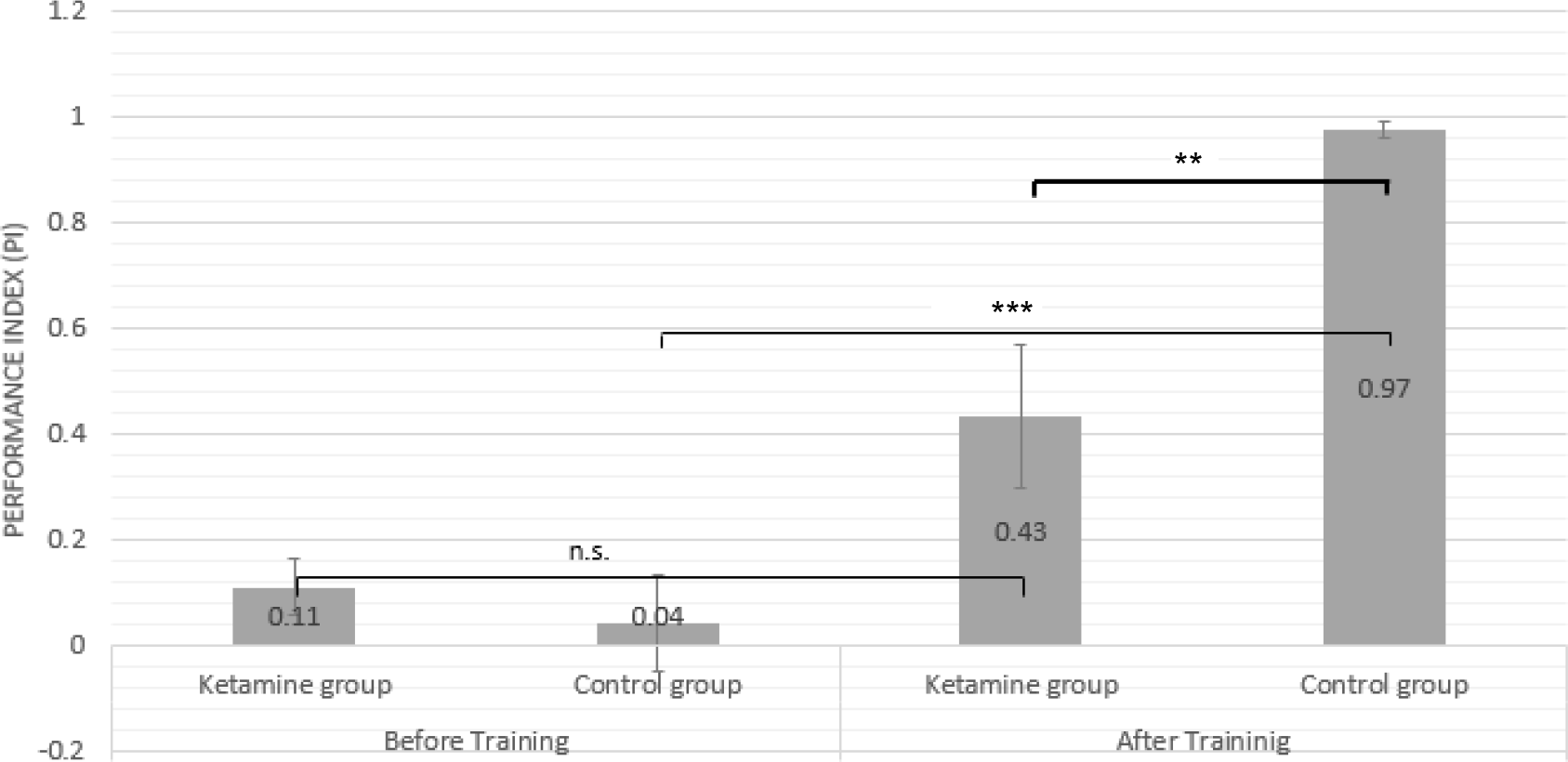
Learning/recall is NMDAR-mediated. Fish in the ketamine group were treated with sub-anesthetic dose of ketamine (2mg/L), a non-competitive glutamatergic antagonist. The treatment lasted for 20 minutes before the training session and continued during the test phase. In the untreated control group (n=6), the PI (mean+/−SE) post-training increased significantly (p<0.001) whereas the increase in PI (mean+/−SE) of ketamine treated group (n=7) was non-significant. Paired (thin line) and unpaired (bold line) t-tests were performed as appropriate to determine the degree of significance. ***p<0.001, **p<0.01

### Cold anesthesia is safe in adult *Danio rerio*

**Figure 4.**
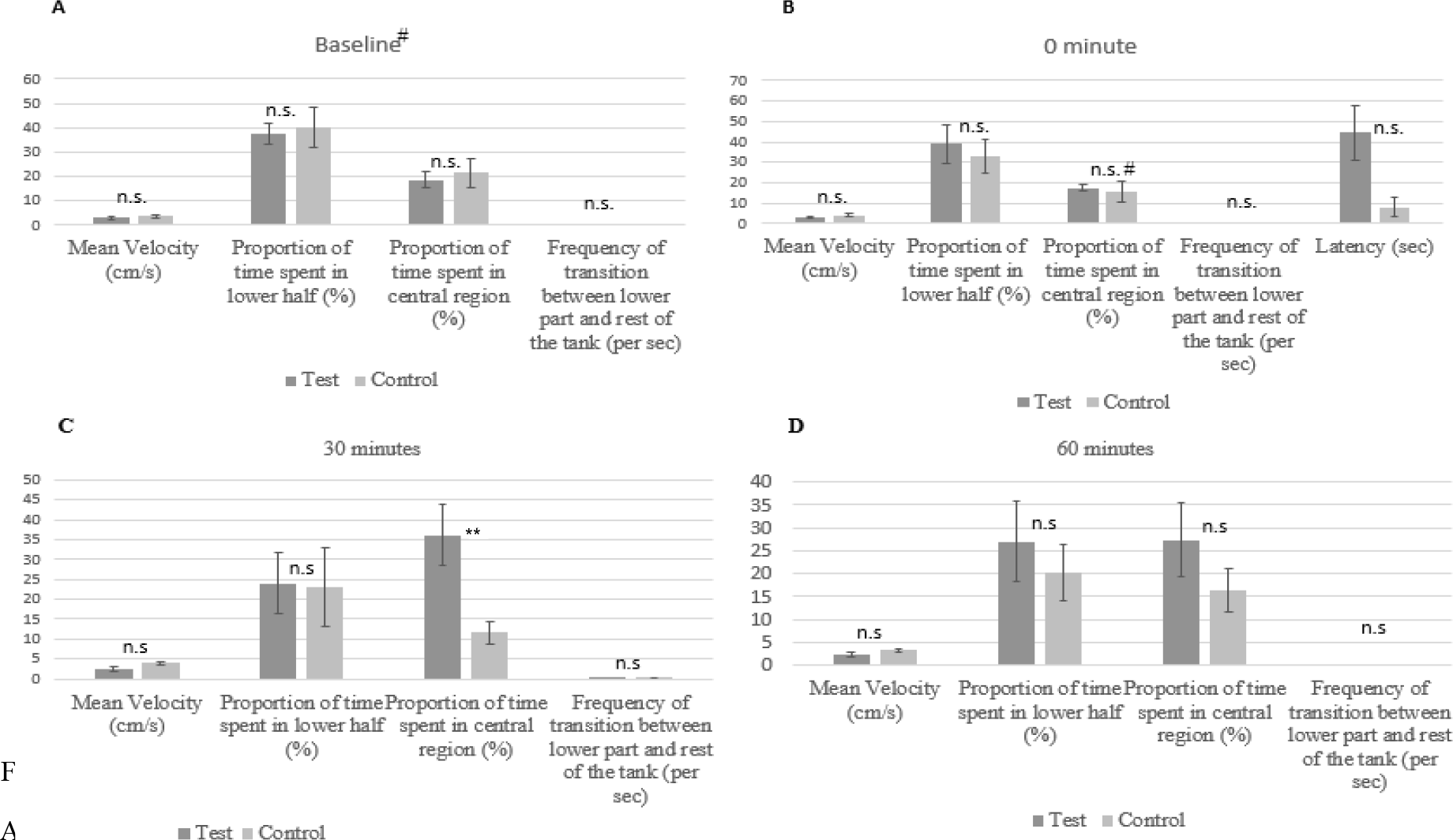
(A to D): Cold anesthesia protocol does not harm swimming behavior in adult *Danio rerio*. A group of fish (n=6) were subjected to the cold anesthesia protocol. Their swimming behavior was recorded at pre-treatment (baseline) and 0, 30 and 60 minutes post-treatment. The recordings were analyzed and the above plotted parameters were quantified (mean+/−SE) to assess the effect of cold anesthesia on the swimming behavior of zebrafish. An untreated control group (n=5) was processed which underwent a similar treatment without the cold anesthesia. (A) In baseline swimming activity, treated and untreated groups do not show any significant difference in any of the parameters. (B) At 0 minute post-treatment, a difference in latency is observed which is still non-significant (p=0.06). (C) At 30 minute, a significant difference is observed only in the proportion of time spent in the central region. (D) By 60 minutes, there are no significant differences in any of the swimming behavior parameters between the treated and untreated groups. As all our tests are performed at 2 hours after the treatment, it is safe to use the cold anesthesia protocol without altering the swimming behavior in the fish. Unpaired t-tests were performed to determine the degree of significance. **p<0.01, #number of control fish in the marked cases were four.

### Post-training hypothermic shock has no effect on memory recall

**Figure 5:**
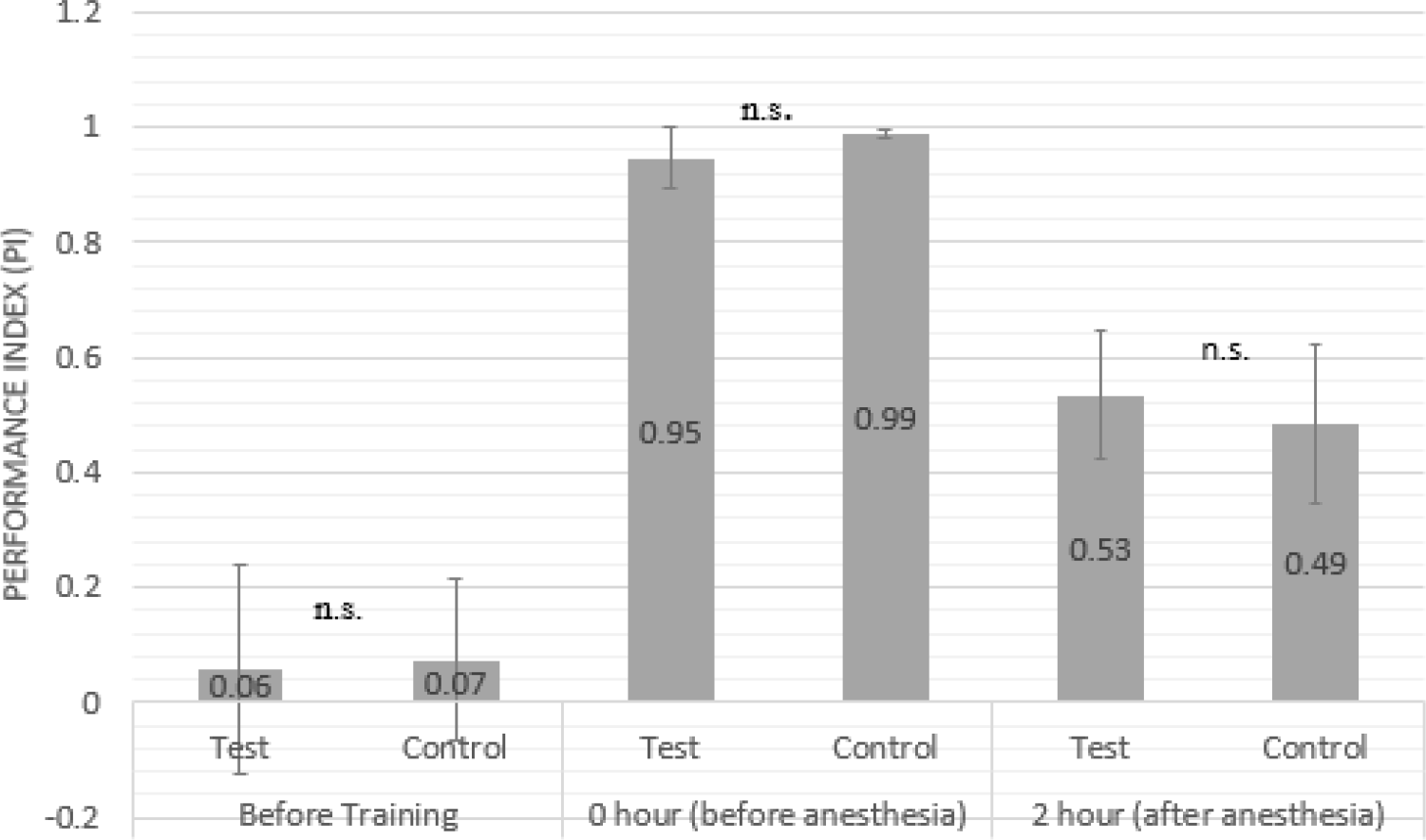
Cold anesthesia post-training does not influence PI. A group of fish was given the cold anesthesia protocol after the training session to study the effect of the cold anesthesia on PI. Mean values are shown on the bar. The differences in the PI between the cold treated test group (n=5) and the untreated control group (n=5) were non-significant, both before the treatment (at 0 minute) and after the treatment (2 hours). Unpaired t-test was performed to determine the degree of significance.

### Post-training tricaine anesthesia has no effect on memory recall

**Figure 6:**
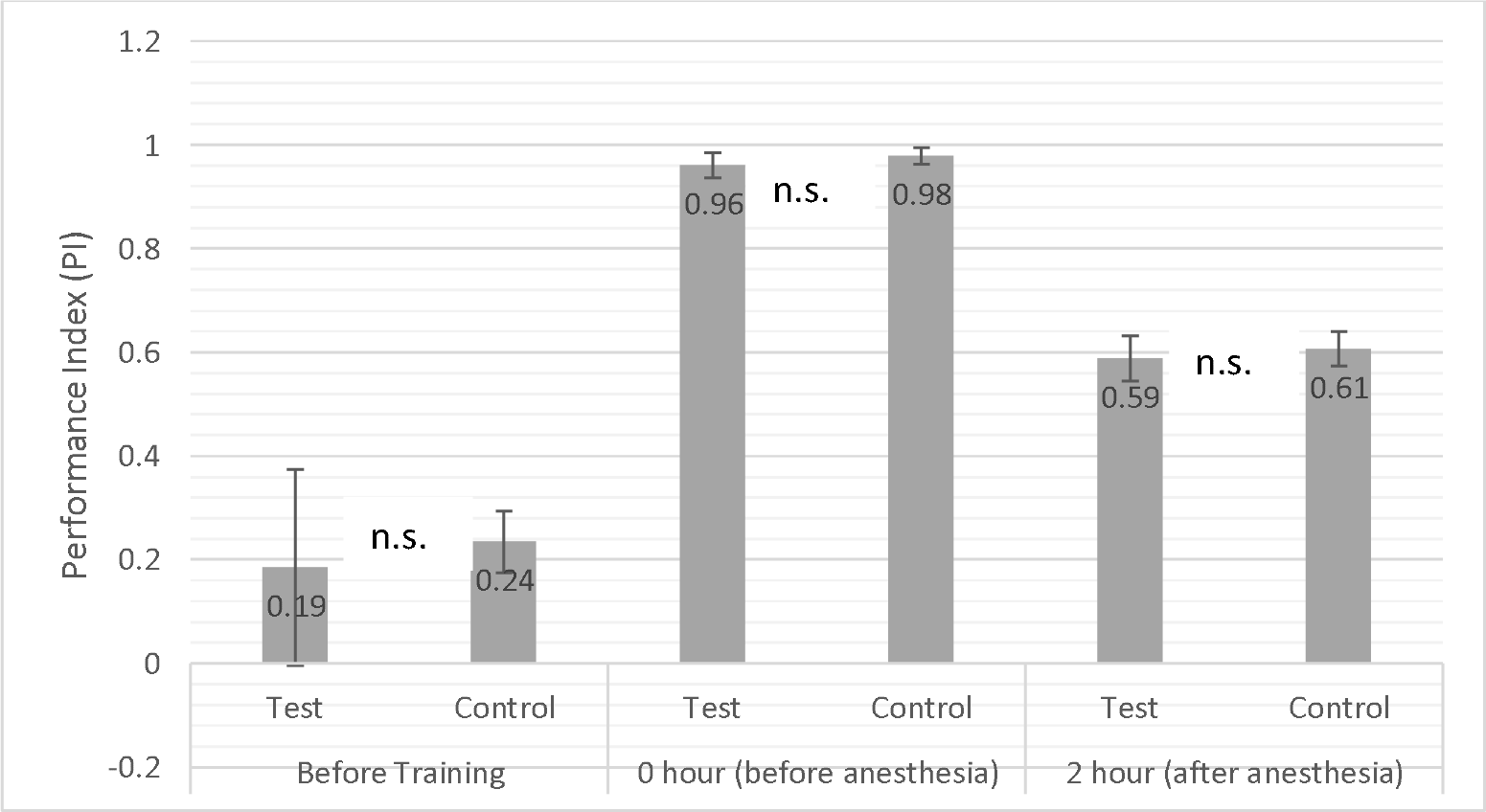
Tricaine anesthesia post-training does not influence PI. A group of fish was given the tricaine anesthesia protocol after the training session to study the effect of the tricaine induced anesthesia on PI. Mean values are shown on the bar. The differences in the PI between the tricaine treated test group (n=3) and the untreated control group (n=4) were non-significant, both before the treatment (at 0 minute) and after the treatment (2 hours). Unpaired t-test was performed to determine the degree of significance.

## DISCUSSION

We used a massed training classical conditioning protocol using a green light at the bottom of the tank as the conditioned stimulus (CS) and electric shock as the unconditioned stimulus (US). Zebrafish showed remarkably robust learning in the fear conditioning assay. The performance index immediately after training was 100%, and the 3 hour memory retention remained significantly higher in the trained group (Figure 1). We carried out the memory tests at various time intervals (0, 3, 6, 12 and 24 hours) in a group of trained fish to capture the rate of memory decay in adult zebrafish on this learning task. We observed a declining trend in the performance index over time, indicating a decay in memory over 24 hours (Figure 2). Imitation learning or conflict inhibition was avoided by carrying out the training and testing in isolation (Gleason and Webber, 1977).

A massed training protocol induces a shorter lasting memory whereas a spaced-training protocol induces a long-lasting protein-synthesis dependent memory. We developed a massed training protocol as we are interested to look at only the short-term memory and its dependence on an anesthesia-sensitive pathway. A test period was selected 2 hours post-training to evaluate memory performance because we had already shown that the performance in trained group, 3 hours post-training, remained significantly higher compared to the control group (Figure 1).

Later, we used ketamine, a non-competitive glutamatergic antagonist, at a sub-anesthetic dose (2mg/L) to understand the learning mechanism at a molecular level. At this concentration, ketamine does not impair swimming or shoaling behavior in adult zebrafish (Riehl *et al*., 2011). We found that in the presence of ketamine, the learning was significantly reduced (Figure 3). The control group in the experiment was an animal group trained in the absence of any pharmacological agent. The test in these animals was done immediately after training (not after 2 hours) to evaluate whether there is any learning at all. The significant decrease in learning in the ketamine-treated group clearly points to a critical role played by NMDA receptors. We are unable to say precisely whether NMDA-R plays a role in learning, recall or both as the animals in our experimental set-up were exposed to ketamine during both, training and testing (recall).

To experimentally evaluate the effect of cold-anesthesia on memory, we had to firstly develop a cold anesthesia protocol which should be safe to use in adult zebrafish without altering its native swimming behavior. A fish group was tested on the cold-anesthesia protocol (see Methods) and its behavior was analyzed at various time intervals, to comment on the effect of cold-anesthesia on swimming behavior. The rationale for choosing the parameters are: mean velocity (a measure of sedation), total time spent in the lower third of the tank (a measure of anxiety in novel tank diving test), cumulative time spent in the central part of the tank (thigmotaxis as a measure of anxiety), initial latency to leave the lower one third of the tank (a measure of sedation and anxiety) and frequency of zone transition between the lower third and the rest of the tank (a measure of anxiety in novel tank diving test). The parameters for the test were adapted from Nordgreen *et al*. (2014). We found that the differences between the cold-anesthesia treated and the control group were non-significant across all the above mentioned parameters at the 60^th^ minute post-training. This was used as a confirmation that the anesthesia protocol was safe to be used in our study as our test experiments were carried out 120 minutes post-training. Although, we had expected the latency parameter at 0 minute to be significantly lower in the treated group, even this turned out be statistically non-significant (unpaired t-test, p= 0.06).

In our experiment to test the effect of cold anesthesia on memory, we employed cold-anesthesia treatment post-training and evaluated the effect on memory by testing the performance at 2 hours post-training. The difference between the performance index of the test and control group was non-significant (Figure 5). When we view this result with the knowledge of *Drosophilla* literature, there are many possible explanations for this observation: (1) The Anesthesia Sensitive Memory (ASM) pathway which is sensitive to hypothermic shock in *Drosophila* may be completely absent in adult zebrafish, at least in a paradigm involving aversive learning. (2) Despite being an ectothermic animal like *Drosophila,* the absence of memory sensitivity to hypothermic shock may indicate that zebrafish have evolved strategies to cope with the physiological damage brought about by the cold-shock. (3) While a possibility is that the ASM is completely absent, the other possibility is that the ARM pathway is simultaneously active and is a significantly larger contributor in the observed memory. This would consistent with the existence of parallel system for memory traces of different kinds in drosophila (Bouzaiane *et al* 2015) (4) It is also possible that, unlike *Drosophila*, the acquisition of short-term memory in *Danio rerio* takes place near-simultaneously with the training which leaves little, if any, time interval to disrupt the process with a hypothermic shock. (It is also important to note that the transfer of the fish from the training set-up to anesthesia set-up in both the cases had a delay of <4 mins.) (5) Of course, another possibility is that the intensity of hypothermic shock that is delivered in the present study is not high enough to disrupt the acquisition of short-term memory.

Although we have not found many studies in the mice literature which have looked at the effect of hypothermic shock post-training on memory retention, a study by Nagy *et al*. (1976) is of particular interest. The authors show that a hypothermic shock delivered after training in infant mice disrupts memory retention when tested at 24hr post-training but has no effect on the memory retention at 1hr post-training. This is similar to our observation in zebrafish for the short-term memory lasting a few hours post-training. Whether the acquisition for short-term memory proceeds rapidly near-simultaneously with the training or it is completely independent of anesthesia, it appears that the mechanisms underlying formation of short-term memories might be more similar in mice and zebrafish when compared to *Drosophila*.

In addition, albeit with a small sample size, we also tested the effect of chemically-induced anesthesia (using tricaine) on this short-term memory trace and again found the memory to be robustly maintained in the test animals after tricaine treatment (Figure 6). Tricaine acts on voltage gated Na^2+^ channels on the neurons (Attili *et al*., 2014). The retention of memory after such a pan-neuronal shut-down of sodium channels post-training also points to the fact that the acquisition of short-term memory may not be critically dependent on the brain activity post-training in *Danio rerio*. Whether, as in mice, this perturbation has an effect on the memory recall at a much later time point post-training is yet to be seen in *Danio rerio*. This has to be addressed using a long-term memory assay.

It is very early to comment on memory pathways in zebrafish that are equivalent to the observed pathways in *Drosophila.* However, it would be relevant and interesting to test how zebrafish mutants in the orthologous genes of *rutabaga* and *radish* (of *Drosophila*) would perform on the learning assay. We may have to specifically direct the mutations to subsystems known to be involved in encoding fear memories in zebrafish. Such tightly regulated spatial targeting can be achieved by driving cas9 using Gal4/UAS system on available tissue-specific Gal4 lines (Donato *et al*., 2016). This will let us further resolve the underlying memory pathways in zebrafish from an anatomical and molecular perspective.

Most studies in zebrafish behavior so far have focused on sensory and motor functions in larval zebrafish, which are more malleable to optical and genetic tools. However, the young larvae do not perform well on learning and memory paradigms. In future, as studies in zebrafish model extend to understand the genetic and neural correlates of learning and memory in adult fish, our study can provide an insight on the acquisition of short-term memory in adult zebrafish.

## Supporting information

Behavior example movie

