## Supplementary material for "An NMDA-R mediated short-term memory resistant to anesthesia in adult *Danio rerio*"

**Massed, simultaneous-conditioning protocol**

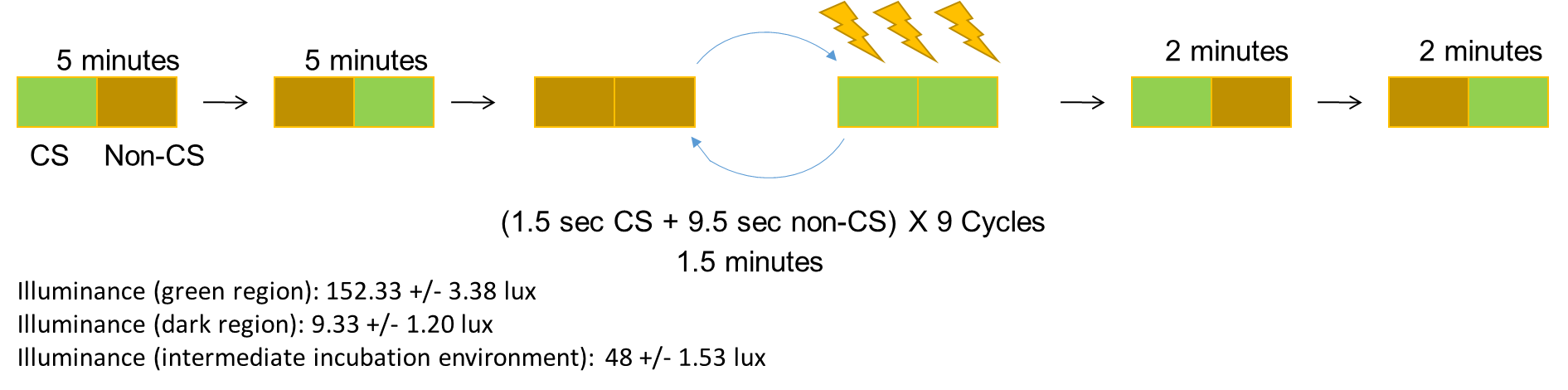

**Representative image showing swimming trajectory of the fish after recovery from cold anesthesia (temperature)**

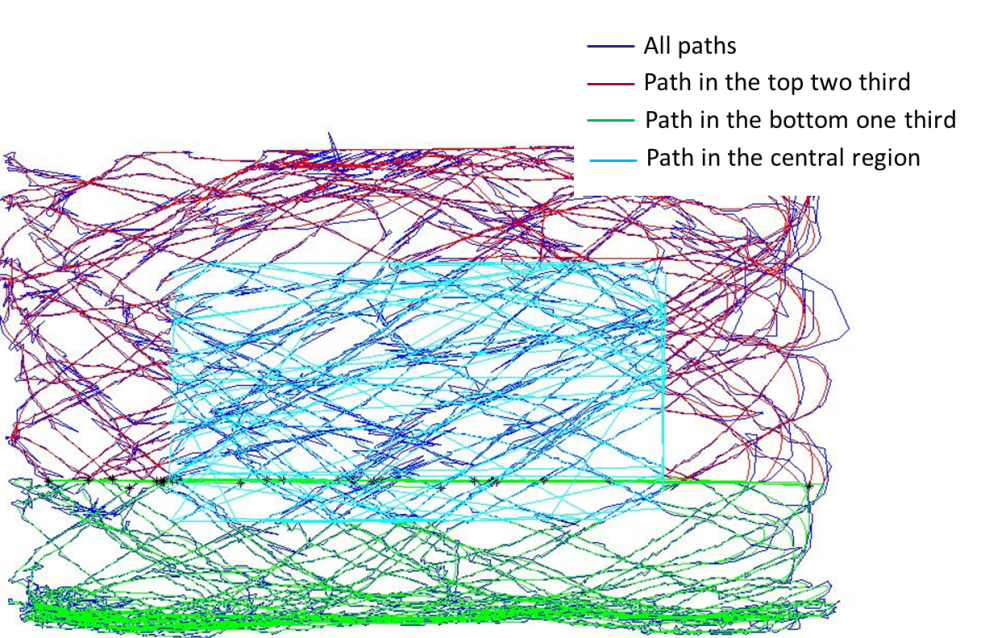

Training setup

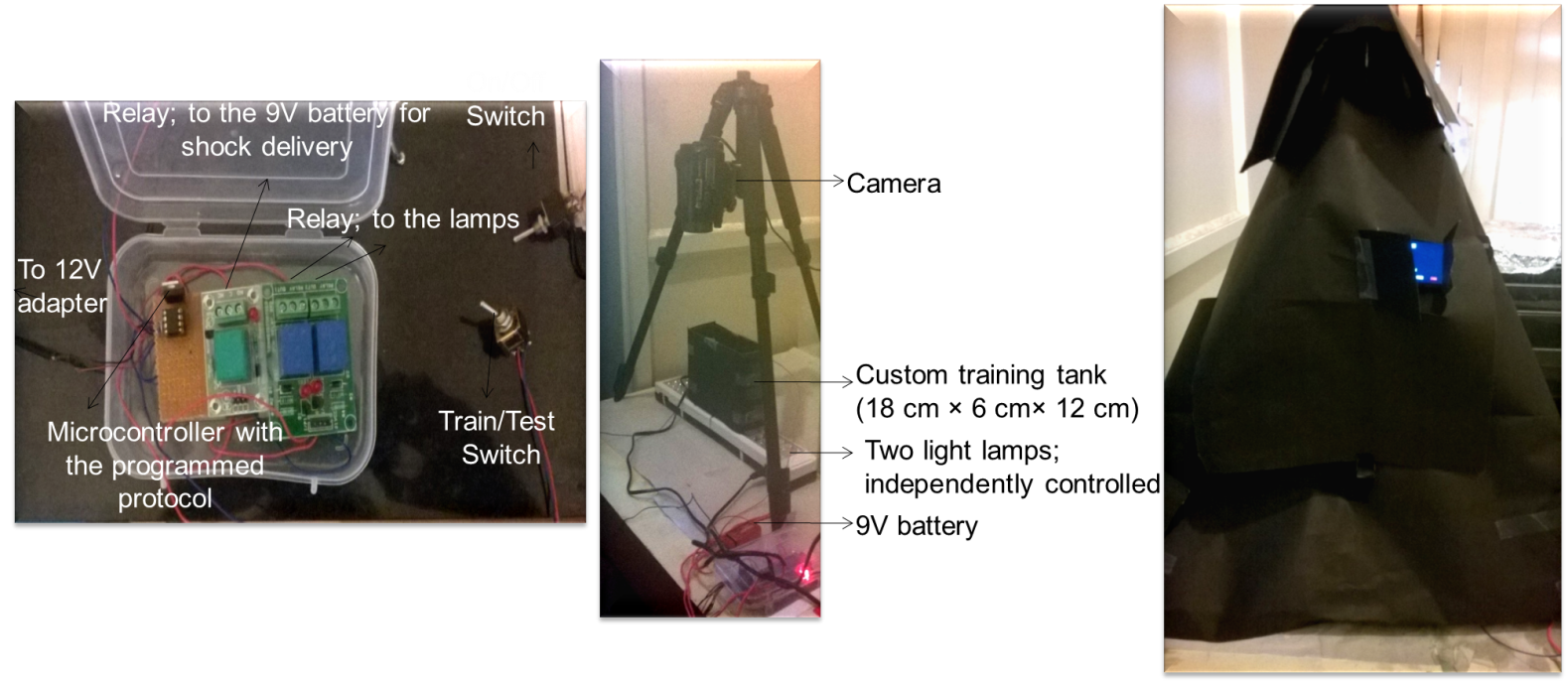
